# A comparative analysis of automated MRI brain segmentation in a large longitudinal dataset: Freesurfer vs. FSL

**DOI:** 10.1101/2020.08.13.249474

**Authors:** Javier Quilis-Sancho, Miguel A. Fernandez-Blazquez, J Gomez-Ramirez

**Affiliations:** Instituto de Salud Carlos III, Fundación CIEN

## Abstract

The study of brain volumetry and morphology of the different brain structures can determine the diagnosis of an existing disease, quantify its prognosis or even help to identify an early detection of dementia. Manual segmentation is an extremely time consuming task and automated methods are thus, gaining importance as clinical tool for diagnosis. In the last few years, AI-based segmentation has delivered, in some cases, superior results than manual segmentation, in both time and accuracy. In this study we aim at performing a comparative analysis of automated brain segmentation. In order to test the performance of automated segmentation methods, the two most commonly used software libraries for brain segmentation Freesurfer and FSL, were put to work in each of the 4028 MRIs available in the study. We find a lack of linear correlation between the segmentation results obtained from Freesurfer and FSL. On the other hand. Freesurfer volume estimates of subcortical brain structures tends to be larger than FSL estimates of same areas. The study builds on an uniquely large, longitudinal dataset of over 4,000 MRIs, all performed with identical equipment to help researchers understand what to expect from fully automated segmentation procedures.

## I. Introduction

Both healthy aging and neurodegenerative and psychiatric disorders experience changes in the structure of the brain [Gado et al., 1983, Snowdon, 2003, Carlson et al., 2008, Fjell et al., 2009]. These changes may include differences of volume, in the morphology of subcortical structures or alterations in the thickness and morphology of the cortex [Yang et al., 2019, Azevedo et al., 2019, Vollmer et al., 2015].

Brain atrophy, either the organ’s volume or specific subfields, can work as a biomarker to reveal the existence of neurological disorderseases, such as Alzheimer’s disease [Pini et al., 2016], Parkinson’s disease [Zhou et al., 2020] and multiple sclerosis [Losseff et al., 1996, Fox et al., 2000, Andravizou et al., 2019] among others. Two well-established markers of dementia are loss of gray matter volume [Beyer et al., 2007, Peter et al., 2014] and the shrinking of the hippocampus [Wang et al., 2003, de Flores et al., 2015]. Being able to detect these signals before behavioral deterioration begins is crucial for the early diagnosis of the disease.

In the era of big data, we have at our disposal an extremely huge amount of datasets that are being fed at ever increasing frequency from sensors and also researchers’ across different scientific fields. Neuroscience deserves a particular mention as one of the most data-rich scientific endeavours. Repositories with thousands of MRI scans freely accessible are already available [Jack Jr et al., 2008, Miller et al., 2016], with more, larger and more complex repositories coming up in the near future.

The main premise of big data applied to brain imaging research is that given enough data, we may be able to characterize brain atrophy and fit the data to a model that makes predictions about the dynamics of the atrophy, that is, not only identify atrophy but its progression in individual basis.

Manual segmentation of the brain is a very time consuming and prone to error task [Collier et al., 2003, Despotović et al., 2015]. To put this in perspective, only to extract the volume of the hippocampus, an expert requires at least 30-40 minutes [Firbank et al., 2008]. Therefore, an exhaustive segmentation of the whole brain could take hours or days of human work which is also prone to errors due to fatigue [Firbank et al., 2008, Starmans et al., 2020]. Hence, the use of manual segmentation on a medium (hundreds of brains) or large scale (thousands of brains) is not an option. For this reason, in this paper we are putting in value the role of automatic procedures of brain segmentation with a large longitudinal dataset. The main advantages of automatic procedures are two: the lack of bias that may have manual segmentation if different experts do the segmentation and the time saving. The goal of this work is thus, to present the current capabilities and promise of automatic procedures for brain segmentation by means of showing performing automatic brain segmentation systematically in a large longitudinal study of 4028 MRI scans collected in a 5 year time span.

## II. Methodology

### Participants

The dataset for this work was collected by a longitudinal study about healthy aging based in Madrid, Spain [Gómez-Ramírez et al., 2019], [Fernández-Blázquez et al., 2020] conceived to investigate the effect of aging on the human brain. The dataset is part of single-center, observational, longitudinal cohort study that started in 2011 and it is still ongoing. The participants were recruited between 2011 and 2013 in Madrid, the cohort started with 990 home dwelling volunteers, between 70 and 85 years old, without any relevant psychiatric, neurological or systemic disorder. Since their inclusion in the study, the volunteers undergo a yearly systematic clinical evaluation which includes medical history, neurological and neuropsychological exam, blood collection and brain MRI. Each subject is examined yearly and information regarding a wide range of factors such as MRI, genetic, demographic, socioeconomic, cognitive performance, subjective cognitive decline, neuropsychiatric disorders, cardiovascular, sleep, diet, physical exercise and self assessed quality of life is collected. The subjects are clinically diagnosed as healthy, mild cognitive impairment or dementia.

The 990 subjects in the first year of the study have the following characteristics: Age 74.72 ± 3.86; 40% Male; 16% heterozygous for the *ϵ*4 genotype of APOE and 0.7% homozygous for the *ϵ*4 genotype of APOE; 17% unschooled, 26% with primary school, 24% with medium or high school and 23% with university studies.

When a subject is diagnosed with dementia, he or she abandons the study, the subjects are free to leave the study by personal choice at any time. Because of that, the number of subjects has been reduced through the years.

### Imaging Study

A total of 4028 MRIs were collected in a 5 year-period, 990 in the first visit, 768 in the second, 723 in the third, 634 in the forth, 542 in the fifth and 371 in the sixth year.

The imaging data were acquired in the sagittal plane on a 3T General Electric scanner (GE Milwaukee) utilizing the following T1-weighted inversion recovery, supine position, flip angle 12∘, 3-D pulse sequence: echo time *Min. full*, time inversion 600 ms, Receiver Bandwidth 19.23 kHz, field of view = 24.0 cm, slice thickness 1 mm, Freq. *×* Phase 288 *×* 288.

In this study we analyze the totality of the 4,028 MRI collected. An amount of data of this size and complexity can not be analyzed by hand. Manual segmentation would be an extremely time consuming job and we do not have at our disposal the resources needed to fulfill this task. For this reason, we examined the performance of automatic procedures to segment the brain in order to obtain the volume and shape estimated of subcortical brain structures.

In the era of big data, automated segmentation is called to play a preponderant role in medical imaging. Automated procedures will be an essential element in the toolkit of radiologists and other health professionals working on medical imaging [Hosny et al., 2018, Thrall et al., 2018]. The reproducibility and time-saving characteristics of automated segmentation are some of the strongest points of AI in the field of imaging. Also, this kind of methods are able to give, not only qualitative, but also quantitative results of great precision and accuracy. However, the field is still in its adolescence and, AI cannot, so far, replace an expert eye in every domain [Topol, 2019]. Nevertheless, the unique capabilities of AI can provide a formidable companion to the expert physician, in particular radiologists, dermatologists and pathologists, all three information-based medicinal practices. Therefore, the characterization of performance of these automated procedures in large datasets as the one presented here, is a matter of great interest now and for the immediate future.

The preproccessing of MRI 3 Tesla images in this study consisted of generating an isotropic brain image with non-brain tissue removed. We used the initial, preprocessing step in the two computational segmentation tool used in this study: FSL pipeline (fsl-anat [FSL tool for processing anatomical images, 2017 and the Freesurfer (recon-all [FreeSurfer cortical reconstruction and parcellation process., 2017]) pipeline.

The stages in the FSL pipeline (in order) are: reorient the images to the standard (MNI) orientation, automatically crop the image, bias-field correction (RF/B1-inhomogeneity-correction), registration to standard space (linear and non-linear), brain-extraction, tissue-type segmentation and subcortical structure segmentation.

The stages in the Freesurfer pipeline (in order) are: surface-based stream, with skull-stripping cerebellum and brain stern removal, two hemispheres separation and brain voxels classification (white matter, gray matter and CSF), and finally brain segmentation (cortical and subcortical).

We run both pipelines in an identical computational setting: Operating System Mac OS X, product version 10.14.5 and build version 18F132. The version of Freesurfer is freesurfer-darwin-OSX-ElCapitan-dev-20190328-6241d26. The version of the BET tool for FSL is v2.1 - FMRIB Analysis Group, Oxford and the FIRST tool version is 6.0.

### Freesurfer and FSL pipelines

The two software libraries for brain segmentation used in in this work are FSL [Woolrich et al., 2009, Smith et al., 2004, Jenkinson et al., 2012] and Freesurfer [Dale and Sereno, 1993, Dale et al., 1999]. Here we will focus on the analysis of structural MRI by the two platforms. In this section we will go through both procedures, showing the main similarities and differences.

FSL [Woolrich et al., 2009, Smith et al., 2004, Jenkinson et al., 2012] allows the processing of structural MRI, functional MRI, diffusion MRI and perfusion MRI. As said, we are focusing on structural MRI. The main tools for FSL based analysis are the Brain Extraction Tool (BET), the FMRIB’s Automated Segmentation Tool (FAST) and the FMRIB’s Integrated Registration and Segmentation Tool (FIRST). The BET [Smith, 2002, Simpson et al., 2012] routine separates brain tissue from non-brain tissue. The FAST [Zhang et al., 2001] routine is able to classify the voxels of the brain into white matter, gray matter and CSF. And the FIRST [Patenaude et al., 2011] uses a Bayesian model to segment the different subcortical structures of the brain. The model sample includes both normal and pathological brains (including cases of schizophrenia and Alzheimer’s disease) and the age range goes from 4 years to 87 [Patenaude et al., 2011]. FSL provides also a tool called Structural Image Evaluation, using Normalisation of Atrophy (SIENA/X) [Smith et al., 2002] that allows the analysis of brain change in different time points. Thus, FSL makes use of the above mention tools i.) to estimate the total brain volume, once bone, fat, muscle and other organs are removed (BET) i.) to perform tissue-based segmentation that differentiates among white matter, gray matter and Cerebral Spinal Fluid (FAST) iii.) to segment putamen, thalamus, amygdala, caudate, pallidum, hippocampus, accumbens and brain stern (FIRST) and finally iv.) to estimate the brain atrophy at two different time points (SIENA/X).

Freesurfer includes tools for processing structural MRI, functional MRI, diffusion MRI and PET data. Again, we are focusing on structural MRI. The cross-sectional analysis starts with the surface-based stream [Dale et al., 1999, Fischl et al., 1999], where the skull is stripped, the cerebellum and brain stern are removed, the two hemispheres are separated and brain voxels are classified as white matter or grey matter and CSF[Fischl et al., 2002, Fischl et al., 2004]. Then, the volume-based stream segments the different subcortical structures of the brain using a subject-independent probabilistic atlas. The training set consists on 40 MRIs, spread in age (10 healthy young subjects, 10 healthy middle-age subjects and 10 healthy elderly subjects) and including pathological brains (10 subjects with AD) [Desikan et al., 2006]. And, finally, the pipeline segments the cortex surface [Fischl et al., 2004]. It uses three different atlas to give tree different cortical parcellation: the Desikan-Killiany atlas [Desikan et al., 2006], the DKT atlas [Klein and Tourville, 2012] and the Destrieux atlas [Fischl et al., 2004, Destrieux et al., 2010]. This pipeline provides the total brain volume, white matter, gray matter, CSF, segmentation of subcortical structures and, unlike FSL, cortical parcellation which is not provided in FSL. Also, the subcortical segmentation is more extensive in FreeSurfer than in FSL because apart from the structures in FSL, Freesurfer also segments cerebellum, optic chiasm, ventricles, vessel and choroid plexus. Freesurfer also has a longitudinal protocol [Reuter et al., 2010, Reuter and Fischl, 2011] which analyzes different MRIs from the same subject across time and reduces the variability between different time points.

Once the general features of each library are shown, it is worth pointing out that Freesurfer has a more extensive subcortical segmentation and also parcels the cortex. In the case of the longitudinal analysis the differences between the two solutions are more prominent. FSL just calculates the whole-brain atrophy between two time points, while Freesurfer is able to analyze every time point simultaneously and gives complete longitudinal segmentation and parcellation. The main disadvantage for Freesurfer is that the protocol take more time to run. To put this in perspective, the Freesurfer cross-sectional segmentation could last around 10 times more that the segmentation by FSL.

## III. Results

We have previously reviewed the main features of both libraries, the next step is to evaluate how they perform in a comparative basis using a dataset of 4028 T1-weighted image MRIs. As mentioned before, our final goal is to evaluate the performance of fully automated methods, without manual segmentation. We aim at comparing how FSL and Freesurfer perform in our large, mono-center dataset. Our focus is on the set of the common brain structures analyzed by both Freesurfer and FSL, that is to say, the total brain volume and putamen, thalamus, amygdala, caudate, pallidum, hippocampus and accumbens.

In this analysis we only apply the cross-sectional protocol to all the images, that is, we treat every MRI as independent from the rest. Therefore, different MRIs from the same subject are treated independently. We used the settings by default in both FSL and Freesurfer processing.

The first approach is to look at the volumes of the brain and the different structures obtained in the first visit from every subject in the dataset. For this first analysis we count on 990 images. Figure 1 depicts histograms of the volumetry measured by both libraries for the total brain volume and the different subcortical structures. Motion and other artifacts can result in poor estimates [Kecskemeti and Alexander, 2020], we excluded 1% of the outliers in each structure, this means that we exclude the 0.5% larger and smaller interval of values.

**Figure 1:**
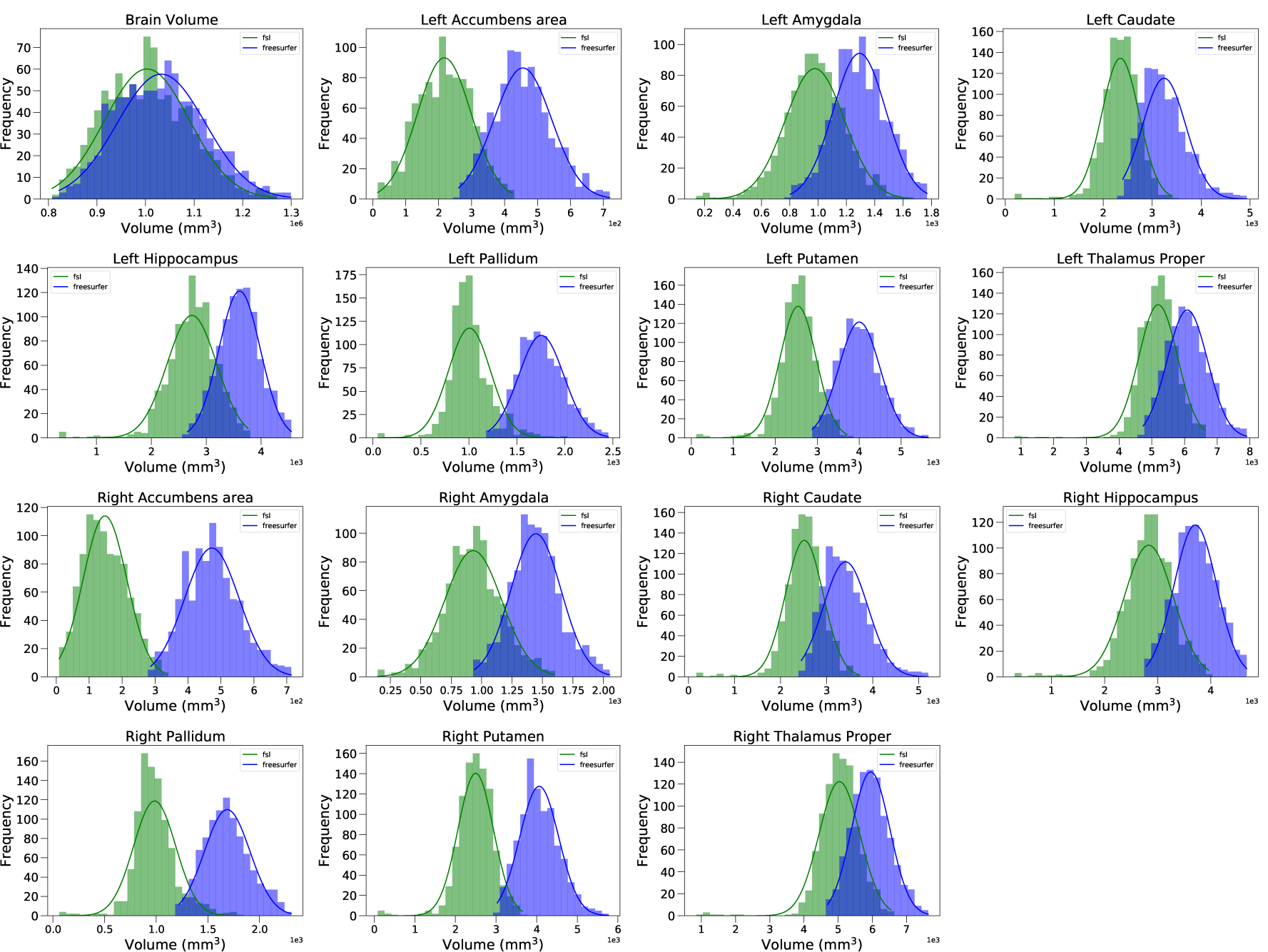
Distributions of every structure analyzed in this study by the two automated methods. The distributions include every resonance obtained in the first visit of every subject (a total of 990). We compare the distributions provided by FSL (in green) and the ones obtained with Freesurfer (in blue). The lines represent the theoretical Gaussian distribution calculated with the mean and the standard deviation of each distribution.

From Figure 1, as expected, if we look at the distributions for each structure we have Gaussian distributions for both FSL and Freesurfer. However, it is worth mentioning that Freesurfer distributions (in blue) have higher values in every structure compared to FSL distributions (in green). This fact is quantified in Table 1 where we can see the means and standard deviations computed by Freesurfer and FSL for every structure. As we can see, the mean of the Gaussian distributions measured by Freesurfer is larger in every structure. Besides, we calculated the p-value comparing both distributions. Although it is straightforward to see in the plots, there is evidence against the null hypothesis -Freesurfer and FSL and distributions are the same-the p-value is lower than 0.01 in every structure. And finally, we calculated the effect size with the Cohen’s *d* to see the standardised difference between the means, observing that with the exception of the brain volume the effect size is very large for every structure.

**Table 1:** Means and standard deviations for the distributions in Figure 1. The last two column shows p-value and Cohen’s *d* from the comparison of the FSL and Freesurfer distributions for each structure. The ∗∗ means that the p-value is lower than 0.01.

| Structure | $\mu_{fsl}$ (mm<sup>3</sup>) | std<sub>fsl</sub> | $\mu_{fr}$ (mm<sup>3</sup>) | std<sub>fr</sub> | p-val | Cohen d (95% CI) |
| --- | --- | --- | --- | --- | --- | --- |
| Brain Volume | 1003131.145 | 87388.925 | 1032458.59 | 91017.684 | ** | 0.33 (0.24-0.42) |
| L Accumbens | 216.333 | 82.316 | 454.939 | 87.938 | ** | 2.80 (2.68-2.93) |
| L Amygdala | 977.378 | 208.738 | 1292.643 | 186.856 | ** | 1.59 (1.49-1.69) |
| L Caudate | 2351.182 | 380.826 | 3244.433 | 444.103 | ** | 2.16 (2.05-2.27) |
| L Hippo. | 2739.324 | 451.388 | 3608.068 | 376.729 | ** | 2.09 (1.98-2.20) |
| L Pallidum | 1005.491 | 220.527 | 1753.616 | 236.138 | ** | 3.27 (3.14-3.41) |
| L Putamen | 2535.503 | 452.667 | 4001.368 | 490.511 | ** | 3.11 (2.98-3.24) |
| L Thalamus | 5194.977 | 594.592 | 6084.683 | 619.566 | ** | 1.47 (1.37-1.56) |
| R Accumbens | 147.22 | 67.014 | 472.016 | 83.022 | ** | 4.31 (4.14-4.47) |
| R Amygdala | 930.895 | 234.438 | 1446.186 | 206.795 | ** | 2.33 (2.22-2.45) |
| R Caudate | 2513.751 | 409.376 | 3410.172 | 485.98 | ** | 2.00 (1.89-2.10) |
| R Hippo. | 2829.544 | 461.866 | 3708.178 | 400.644 | ** | 2.03 (1.92-2.14) |
| R Pallidum | 983.657 | 206.276 | 1687.419 | 221.328 | ** | 3.29 (3.15-3.42) |
| R Putamen | 2496.482 | 443.494 | 4062.842 | 480.476 | ** | 3.39 (3.25-3.53) |
| R Thalamus | 5037.812 | 599.508 | 5952.144 | 559.42 | ** | 1.58 (1.48-1.68) |

So, the statistical tests shown in Table 1 reveals that Freesurfer tends to estimate larger volume values than FSL.

### Self-consistency of the automated methods

Considering the results obtained we might question the consistency of the automated methods. The differences in volume estimates obtained by Freesurfer and FSL might pose a challenge about their consistency in the task at hand, segmenting of MRI scans. However, we need to take into account the experimental design and the characteristics of the collected dataset. As said, the study is longitudinal, however we treated every MRI in our study as independent and the analysis applied was cross-sectional. So, in our dataset we have several images from the same subject at different times (with a time lapse of about 1 year ±2 months between images).

One year is not a long time to expect strongly significant changes in the brain structure and the different volumes of a brain. In these studies [Fox and Freeborough, 1997, O’Brien et al., 2001, Enzinger et al., 2005] the authors show that the mean change in volume that a whole healthy adult brain experiences in a year is lower than 0.5%. While the studies in [Fjell et al., 2009, Fjell et al., 2013] do the same analysis but focusing on the subcortical structures and the differences in volume measured are lower than 5% with means lower than 1% of shrinking for every structure in healthy aging. Hence, a sensible way to test the self-consistency of the codes might be to measure the differences from the same subject between two consecutive years. We would expect, according to the literature, small changes between the images for same subject in two time points spanned one year time.

Following this argument, we calculated the increase/decrease of the volume of the whole brain and the different subcortical structures (in percentage) for every subject between every two consecutive years. This means that if we have a subject with *n* consecutive MRIs, we calculated

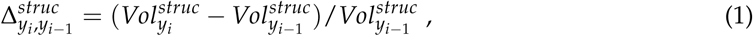

where 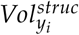 is the volume of a structure, *struc*, for a particular year, *i*, for all the years from 2 to *n*.

The results from Freesurfer are shown in Fig. 2. To clean the distribution from possible outliers and as mentioned before, we exclude the 0.5% larger and smaller interval of values. Focusing first in the brain volume, we can see that the distribution is centered close to zero but skewed towards negative values. This is as expected, since the pass of time will produce a progressive shrinkage of the brain. However we can see that the decrease is really small. Also the limits of the distribution are small, around [-0.045, 0.025] (min. 0 and max. 1), which is expected as well.

**Figure 2:**
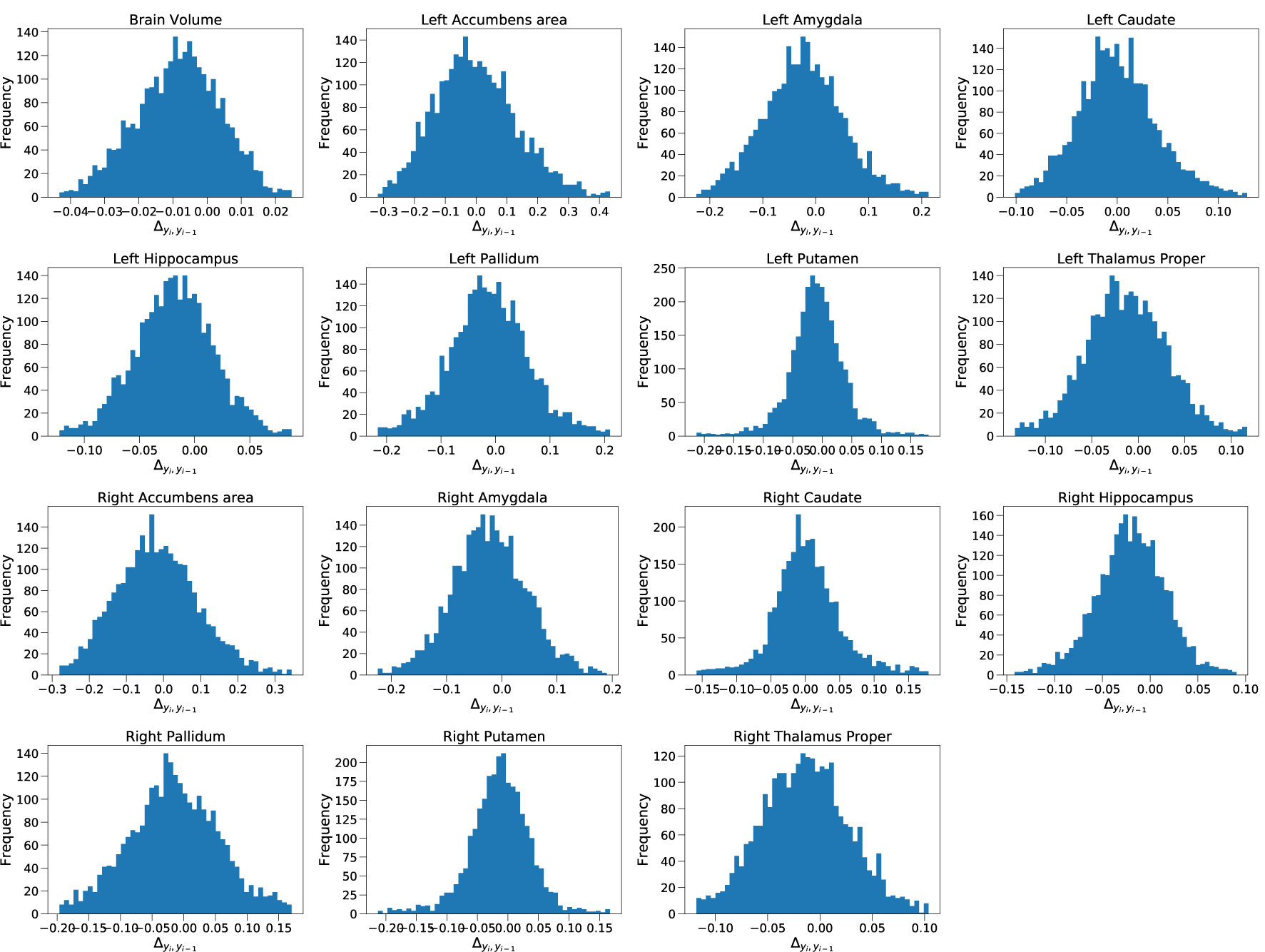
Variations in the volume of the different brain structures between two consecutive years (Eq. 1) measured by Freesurfer.

Looking now at the different structures, we see the same pattern in the brain volume. The distributions are centered around 0 but towards the negative side of the axis. However, the variability is much larger in subcortical structures than it is in the total brain volume. While the maximum change in terms of brain volume is a decrease lower than 0.05, in the case of, for example, the left accumbens area we can see an increase of more than 0.4. This is an extreme case and the differences are below 0.25 (in some case even lower than 0.15) for all the cases from every structure except for the accumbens, which only 62 and 159 cases out of 2743 overcome this 0.25 limit for the right and left part, respectively. The losses are always constrained within the range from 0 (total preservation in between years) to 1 (total loss in between years).

In Figure 3 we show the results from FSL. We have strong similarities with the results obtained from Freesurfer, although for FSL there are more outlier cases discarded. We excluded in this case the 0.025 larger and smaller interval of values (the 100 largest/smallest values), five times more than the Freesurfer case. But as we can see in Figure 3 the resulting cases form a distribution close to a Gaussian.

**Figure 3:**
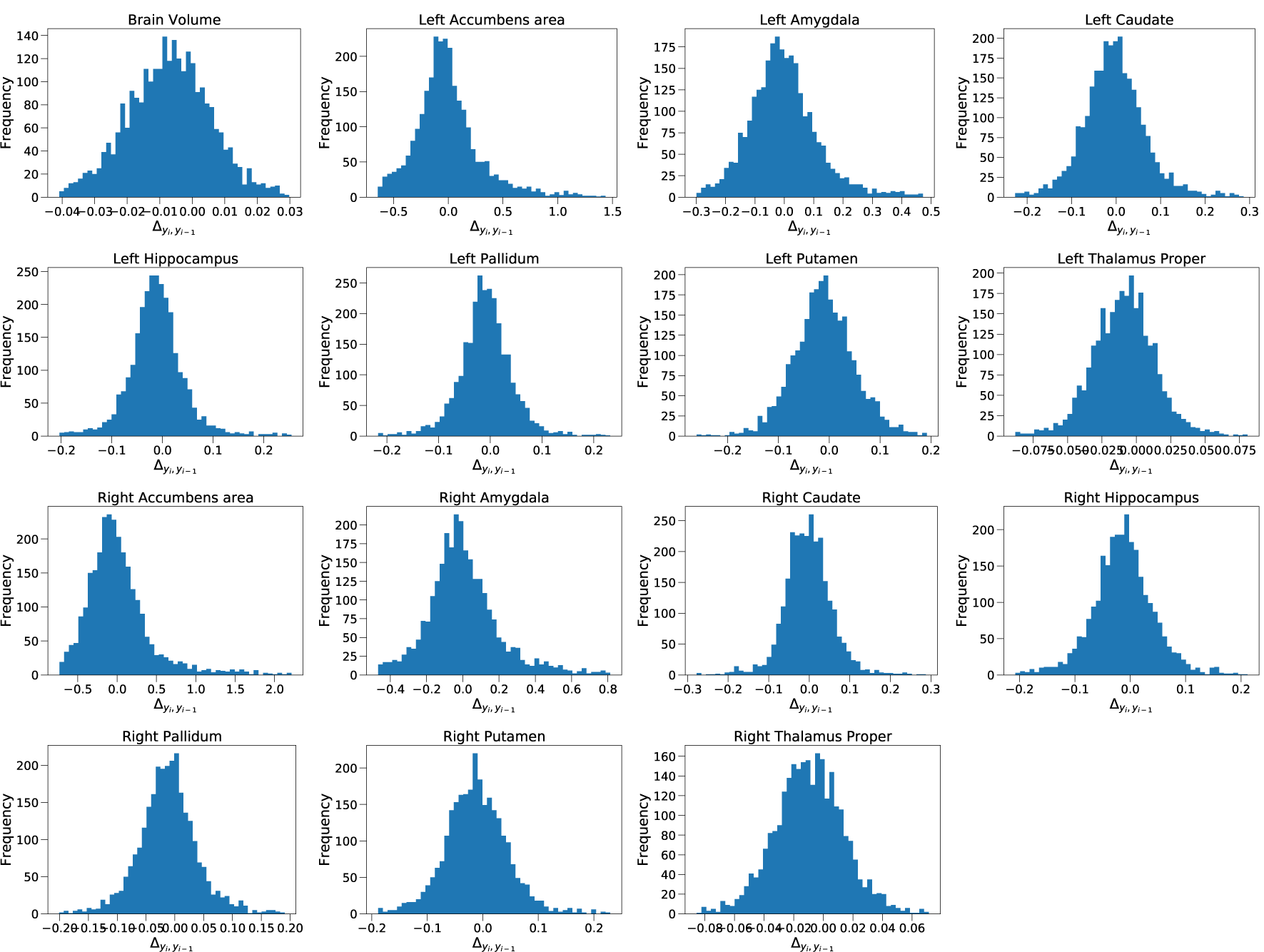
Variations in the volume of the different brain structures between two consecutive years (Eq. 1) measured by FSL.

We start again with the total brain volume. With the outliers removed the changes are lower than a 0.04, which again is what we would expect. Now for the different brain subcortical structures the variability increases again. After removing the outliers the distributions have similar features as in the case of Freesurfer. They also are centered close to zero but skewed towards the left or negative values. The variability is small for the caudate, hippocampus, pallidum, putamen and thalamus, where again almost every case is below the 0.25 difference, or lower. However, the amygdala has cases overcoming the 0.5 limit and specially significant is the accumbens with larger differences. Therefore, for these two structures, amygdala and accumbens, the variability is considerably larger in FSL than in Freesurfer.

In Table 2 we the results exposed in Figures [2 3] exposed in a quantitative way. As we can see, the mean brain shrinking is of 0.007 for FSL and 0.008 for Freesurfer, which is similar to the results obtained in references [Fox and Freeborough, 1997, O’Brien et al., 2001, Enzinger et al., 2005], commented before. Concerning the subcortical structures, all the means are lower than 0.01 except for the right amygdala in both libraries and the right hippocampus in the case of Freesurfer. Focusing first in Freesurfer, only the mean of the caudate (left and right) is larger than zero, while the mean volume of the rest of structures decreases in a year. This result is very similar to the results in References [Fjell et al., 2009, Fjell et al., 2013], commented before, where the mean shrinking of every structure is lower than 1%.

**Table 2:** Means and standard deviations of the relative changes for every structure within a year. The distributions are shown in Figures [2 3].

| Structure | $\mu_{fsl}$ | $std_{fsl}$ | $\mu_{fr}$ | $std_{fr}$ |
| --- | --- | --- | --- | --- |
| Brain Volume | -0.007 | 0.015 | -0.008 | 0.012 |
| Left Accumbens area | 0.018 | 0.414 | -0.001 | 0.133 |
| Left Amygdala | 0.004 | 0.169 | -0.024 | 0.076 |
| Left Caudate | 0.005 | 0.187 | 0.0004 | 0.039 |
| Left Hippocampus | -0.007 | 0.122 | -0.018 | 0.036 |
| Left Pallidum | -0.001 | 0.163 | -0.012 | 0.073 |
| Left Putamen | -0.001 | 0.218 | -0.01 | 0.047 |
| Left Thalamus Proper | -0.008 | 0.061 | -0.013 | 0.044 |
| Right Accumbens area | 0.069 | 0.671 | -0.016 | 0.11 |
| Right Amygdala | 0.02 | 0.291 | -0.022 | 0.067 |
| Right Caudate | 0.005 | 0.18 | 0.003 | 0.05 |
| Right Hippocampus | -0.01 | 0.113 | -0.021 | 0.036 |
| Right Pallidum | -0.004 | 0.17 | -0.016 | 0.07 |
| Right Putamen | 0.006 | 0.315 | -0.014 | 0.05 |
| Right Thalamus Proper | -0.008 | 0.071 | -0.014 | 0.041 |

A key issue to be discussed in this work is the standard deviation, which is much larger than the ones obtained in the references mentioned. The results from FSL, however, are more dissimilar than the ones in the mentioned references. For FSL, the mean volume of half of the structures increases. Nevertheless, it is important to remark that the means is very close to zero and the standard deviation are, again, quite large.

Summarizing, we can conclude that both FSL and Freesurfer are self-consistent. As mentioned, we treated every image as independent. The differences between two images from different years of the same subject tend to be small. As several studies pointed out, the changes in brain volume within a year tend to be neglectful, and finding otherwise in this analysis would make the results untrustworthy. But with the results found, both FSL and Freesurfer seem to be self-consistent. This fact is more strict for Freesurfer, where the number of outliers is much lower and the variability is smaller than FSL.

After checking the self-consistency of both solutions we studied the equivalency between them. To study this, Figure 4 shows the results for Freesurfer and FSL face to face so to speak, for either the brain volume and the subcortical structures. Fig. 4 shows that there is no a linear pattern that relates the FSL results with the Freesurfer ones. We see for every structure a cloud of random points with a variability that coincides, obviously, with the variability seen in Figs. [2, 3].

**Figure 4:**
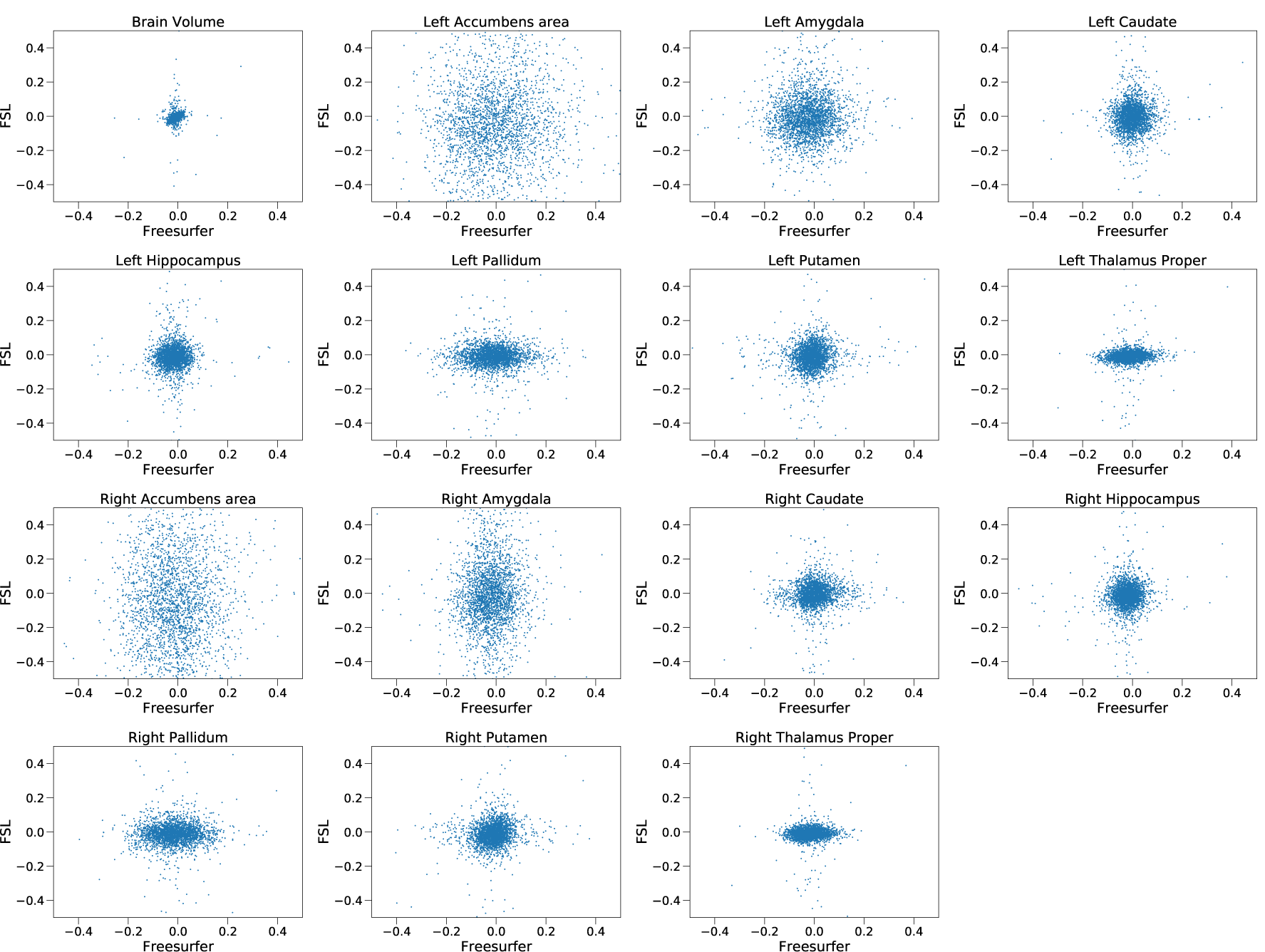
Variations in the volume of the different brain structures between two consecutive years (Eq. 1) confronting the results of Freesurfer and FSL.

If both types of segmentations, FSL based and Freesurfer based, were related we would expect some kind of increasing line or joint pattern. This would imply that a larger measure by FSL would be translated in a larger measure in Freesurfer and viceversa. However we do not observe this. Rather, the lack of structure gives a clear message: there is no apparent relation between the segmentation given by FSL and the one from Freesurfer.

## IV. Discussion and Conclusions

In this study we have studied the results coming from the outcome of applying FSL and Freesurfer to over 4000 MRI dataset of longitudinal data of elderly individuals. The goal was to test the power of automated methods and the advantages that this approach may or may not have.

In Fig. 1 we checked that the distributions of the volumes measured by each library follow a Gaussian distribution. But more important, in Figs. [2, 3] we shown that the changes in volume of the different subjects within a year tend to be small, even when the images are treated as independent. These results are consistent with References [Fjell et al., 2009, Fjell et al., 2013], where the segmentation was made manually. But the main dissimilarity between those studies and this work is the size of the standard deviation. The larger standard deviations obtained with this data imply that automated methods are still not as precise as manual segmentation and the variability is larger.

It is important to remark that by design the protocol followed here is fully automated, without any manual corrections. Manual revisions could reduce the possible errors in the segmentation and the large variability. However, one of the goals of this study is to characterize the performance of FSL and Freesurfer at *face value*. Nevertheless, when dealing with datasets as it is our case of thousands of MRIs, manual correction of each image is not an option. The ever-increasing brain data increase in volume, velocity and complexity calls for pipelines fully automated.

Interestingly, in Fig. 4 we observe a lack of linear relationship between the volumes estimated by FSL and Freesurfer. This result might sound counter-intuitive and deserves and explanation. Since we are analyzing the same MRIs with both libraries, common sense would expect a linear correlation between their results, for example, a large/small estimation by Freesurfer would imply a large/small estimation by FSL as well. Instead, we observe cases with large volumes measured by Freesurfer that have small volumes measured by FSL and the other way around. Nevertheless, it is not the first time this incongruity manifested. In [Schoemaker et al., 2016] differences comparable to ours were exposed in volume estimates of the amygdala and hippocampus in children. In this study both methods overestimated the volumes compared to manual segmentation, but the volumes estimated by Freesurfer were larger than the ones obtained with FSL, as we observed in our study. More similar to our cohort is Rane et al. in [Rane et al., 2017] where they compared both methods applied to different subcortical structures for old adults. The results of this study indicated larger estimations for the caudate, putamen, hippocampus, amygdala and accumbens from Freesurfer in comparison with FSL, as we found in our study. However, the estimations of the pallidum and thalamus were larger for FSL, which contradicts our results. In [Seixas et al., 2010], the authors studied the volume of the hippocampus of healthy adults finding that the measurements from FSL are larger than the ones from Freesurfer, which also is opposed to our results. The hippocampus and amygdala have been studied also in [Morey et al., 2009], and the results are compatible with ours, Freesurfer estimates larger values for the amygdala and the hippocampus values are very similar for both FSL and Freesurfer. The thalamus and the pallidum have been studied in [Makowski et al., 2018] and in both amygdala and the hippocampus, the estimated volumes by Freesurfer were larger than the ones from FSL. And to conclude, in [Morey et al., 2010], the authors found larger volumes measured by Freesurfer in the amygdala but FSL estimated larger volumes for the accumbens, cudate, hippocampus, pallidum, putamen and thalamus.

It is hard to determine the origin of these inconsistencies between studies. The main problem with most of the studies are the modest size dataset, typically less than a hundred subjects. Differences between studies with few resonances could be a statistical effect and it is hard to extract definitive conclusions from them. Also even when using the same library, FSL or Freesurfer, using one version or another or a different operative systems, can provide distinct results [Gronenschild et al., 2012]. However, in our study we are working with 4000 resonances, with the same scanner, same computational setting and same personnel which gives a particular quality to our study vis a vis studies with smaller sample or that combine MRIs from different scanners.

We argue that the main reason for the inconsistencies between the estimates for FSL and Freesurfer might be found in differences in the segmentation protocol. Different boundaries for the structures would translate in different volumes when a particular MRI is analyzed with one library or other. Another factor that can be important is the dataset used to train the segmentation codes. Our dataset is entirely from elderly people. Depending on how represented is this sector of the population in each of library’s brain template, the results could be more/less trustworthy and resulting in differences between FSL vs Freesurfer. Differences in the segmentation protocol or differences in the atlas used to train the codes are likely to affect to the segmentation estimated.

## Acknowledgements

This work was supported by funding *Fundación General de la Universidad de Salamanca, Centro Internacional sobre el Envejecimiento, CENIE*. Program PILEP+90, program grant (0348_*CIE*_6_*E*).

## Ethics declarations

The authors declare no competing interests.

